# CCL5/CCR5 signaling modulates depression-relevant behavior, neuronal oscillations, and long-term depression of synaptic activity

**DOI:** 10.1101/2025.09.12.675856

**Authors:** Katie Hummel, Lara Stefansson, Karli Gilbert, Matthew Amontree, Junfeng Ma, Daniel Pak, Katherine Conant

## Abstract

Major depressive disorder (MDD) is a debilitating disorder, often associated with perseverative thinking and anxiety. Localized reductions in pyramidal cell activity may contribute to associated symptoms, and effective antidepressant treatments typically enhance overall neuronal excitation. CCL5 is a chemokine that has been shown to reduce excitatory-neuronal activity, and is also increased with MDD and conditions that increase MDD risk. Here, we investigate the CCL5/CCR5 axis for its ability to modulate depression-relevant endpoints that are diminished in MDD, including neuronal oscillations, as well as biochemical and behavioral correlates of the disorder. In comparison to wildtype mice, CCR5 knockouts had increased gamma and theta power, and stronger theta/high-gamma phase amplitude coupling during dark-cycle EEG recordings. Compared to strain-matched wildtype mice, CCR5 knockouts also demonstrated reduced anxiety, increased sucrose preference, and improved associative memory. Proteomic analysis of the hippocampus showed that CCR5 knockouts had reduced levels of the GABA receptor alpha-4 subunit, which mediates tonic inhibition and restricts pyramidal cell plasticity. In complementary primary neuronal culture studies, CCL5 diminished GSK-3β activity and impaired NMDA-dependent long-term depression (LTD), a form of plasticity that promotes cognitive flexibility. In addition, CCL5 signaling increased parvalbumin expression in GABAergic neurons through a CCR5-dependent manner. In combination with the ability of CCR5 to restrain gamma oscillation power and LTD, our data raise the possibility that CCL5/CCR5 signaling inhibits neuronal excitation through increased PV+ interneuron activity. Moreover, data are consistent with the possibility that CCR5 antagonists might share the ability of established antidepressants to both increase PC excitation and reduce PC inhibition.

**Significance Statement:** Major depressive disorder (MDD) is a global leading cause of disability, and is associated with increased chemokine activation and inflammation. In this study, we investigate how the CCR5/CCL5 chemokine axis regulates behavioral and cognitive endpoints associated with MDD. This study aims to provide insight to how chemokine signaling underlies mood and behavioral symptoms of neuropsychiatric disorders. We hope this research supports further investigation of CCR5 antagonists for MDD and related mood and anxiety disorders.

## Introduction

Major depressive disorder (MDD) is a common, yet serious condition associated with an increased risk for substance use disorders and suicide. Co-morbid mood and cognitive symptoms may include anxiety, anhedonia, working memory deficits, and inflexible or perseverative thinking. The underlying physiology of the disorder is complex and heterogenous, but generally decreases cortical excitation [1]. Effective anti-depressant treatments can, in part, correct this through directly effecting excitatory glutamatergic neurons, including neurotrophin-induced structural changes, as well as through indirect reductions of pyramidal cell (PC) inhibition [2–5]. Still, an estimated one-third of patients with MDD experience treatment resistance to typical antidepressants, emphasizing the need for novel drugs and adjuvant therapies for pervasive symptoms.

C-C chemokine receptor 5 (CCR5) is a chemo-attractant cytokine receptor that acts as a co-receptor for the human immunodeficiency virus (HIV-1), and is targeted by the highly selective antagonist maraviroc (MVC) for treatment of HIV. CCR5 antagonists are known to attenuate extracellular matrix (ECM) accumulation in peripheral organ systems, and thus have the potential to diminish maladaptive ECM accumulation in MDD [5–9]. Consistent with this, C-C motif chemokine ligand 5 (CCL5) is increased in MDD, and reduced by antidepressant treatment [10–12]. Perhaps more important is work by Silva and colleagues, showing that CCL5 impairs PC excitability and long term potentiation of synaptic transmission (LTP), both of which directly reduce PC excitation [13].

With the goal of determining whether CCR5 antagonists should be considered as an antidepressant or adjunct therapy for MDD, we investigate the role of the CCR5/CCL5 axis in molecular and physiological correlates of mood and cognition. To that end, we use a battery of behavioral assays to assess anxious and anhedonic behavior, as well as working memory, in CCR5 knockout (CCR5 KO) mice and wildtype (WT) controls. We also record *in vivo* continuous local field potentials (LFPs) in order to compare global changes in cortical gamma and theta power, which are altered in MDD [14–16]. To identify novel molecular contributors of mood and memory regulation, we perform an unbiased proteomic analysis of the hippocampus. Finally, we use *in vitro* neuronal culture studies to understand the regulatory role of the CCR5/CCL5 axis in signaling cascades critical for plasticity and memory. CCR5 is a G-protein coupled receptor (GPCR) known to activate downstream G-protein dependent and β-arrestin-dependent signaling cascades, which modulate glycogen synthase kinase-3 beta (GSK-3β) activity. GSK-3β plays a critical role in long term depression of synaptic activity (LTD), so we examine how this plasticity can be modulated by CCL5/CCR5 signaling. Evidence suggests that GSK-3β activity may influence parvalbumin (PV) expression, and so we examine potential effects of CCL5 on the modulation of inhibitory PV+ interneurons.

## Results

### CCR5 deletion increases sucrose preference and hedonic motivation

CCL5/CCR5 inhibition has been shown to increase LTP and pyramidal cell excitability, as do highly effective antidepressant drugs [1]. Consistent with this, CCR5 knockout mice show enhanced cortical LTP, and improved memory [13]. To better understand the effects of CCL5/CCR5 signaling on cognitive and behavioral correlates related to major depressive disorder, we compared strain matched WT and CCR5 KO mice.

To assess anhedonia and motivation, we used the sweet drive test (SDT), a modified sucrose preference test, using normal chow food as well as a sweet, grain-based cereal. After being habituated for 2 days to a modified T-maze apparatus with all boundaries removed, mice were fasted for approximately 12 hours and placed into the center arm of the maze. For 10 minutes, mice were allowed to explore the maze and freely consume the pre-weighed chow and sweet cereal (500 mg each). Mice completed 3 total trials, 48 hours apart to minimize fasting stress. After each trial, the pellet and cereal were weighed to measure consumption (mg). Across 3 trials, CCR5 KO mice showed a higher preference for the sweet cereal (*t*(18) = 2.215, *p* = 0.0399, *d* = 0.991) compared to WT mice (*n* = 10 per group) (Figure 1.A). Although arms were switched for sweet and normal food after every trial, it is of note that by trial 3, CCR5 KO mice had a strong, albeit not significant, preference for entering the sweet arm first, as indicated by the open circles, compared to WT mice, who preferred to enter the chow arm first, as indicated by filled squares.

**Figure 1.**
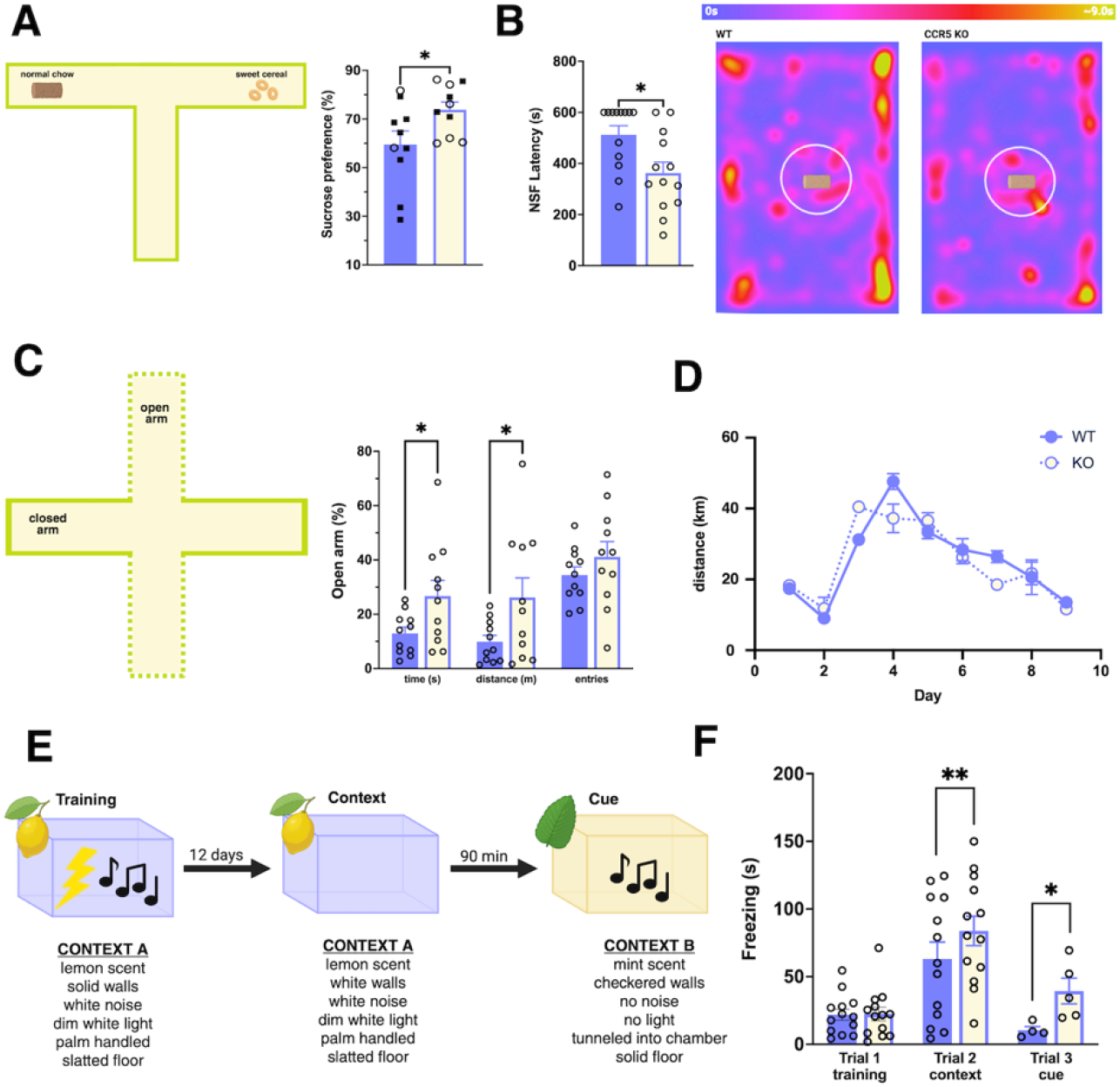
CCR5 KO mice exhibit decreased anxiety and anhedonia-like behaviors, as well as enhanced cue and contextual memory. **(A)** In the sweet drive test (SDT; left), CCR5 KO mice (n = 10) showed an increased sucrose preference compared to WT controls (n = 10 per group, t(18) = 2.215, p = 0.0399, d = 0.991). **(B)** CCR5 KO mice also demonstrated greater motivation and reduced hyponeophagia in the novelty suppressed feeding (NSF) paradigm, where they exhibited a significantly lower latency to feed (n = 12 per group, t(24) = 2.702, p = 0.0125, d = 1.06). Heatmaps (right) show the apparatus setup, as well as mean time in each location, for WT (left) and CCR5 KO (right) groups. **(C)** In the elevated plus maze (EPM; left, n = 11 per group), CCR5 KO mice spent a higher proportion of testing time (seconds) in the open arm (t(20) = 2.217, p = 0.0384, d = 0.945). Distance, which was calculated as (open arm meters / total meters), was also significantly different between groups (t(20) = 2.162, p = 0.0429, d = 0.921). Percent of entries to the open arm were not significantly different (t(20) = 1.503, p = 0.3048, d = 0.4487). **(D)** Behavioral differences were not attributed to increased locomotion, as AUC analysis were no significant locomotor differences between genotypes over 9 days (n = 12 mice per group; t(22) = 0.1902, p = 0.8509, d = 0.077). Each dot represents the daily group average. **(E)** The delay-cued/contextual fear conditioning paradigm consists of 3 trials. Trial 1 is comprised of a habituation phase in context A, followed by a training phase where the animal receives an unconditioned stimulus (mild foot shock) in combination with a conditioned stimulus (tone). After 12 days, the animal completes 2 more trials. For trial 2, the animal is placed back into context A to measure contextual learning. 90 minutes later, the animal is placed into context B with the conditioned stimulus, to measure cue learning. Trial 3 consists of a secondary phase where the cue is terminated and the animal is allowed to freely explore context B, to measure fear generalization. **(F)** In a 2-way ANOVA of z-normalized freeze time, CCR5 KO mice showed stronger associated learning in both the cued (q(22.35) = 4.315, p = 0.0058, d = 1.137) and contextual (q(4.625) = 4.168, p = 0.0352, d = 0.489) learning trials. There were no significant changes in baseline (q(23.02) = 0.1679, p = 0.9065, d = 0.0319) or generalization (q(5.999) = 1.804, p = 0.2492, d = 0.825) measures. CCR5 KO mice showed a higher disparity between tone and no tone in context B versus wildtype (CCR5 KO n = 5, q(7) = 2.331, p = 0.0526, d = 0.703; WT n = 4, q(7) = 0.2044, p = 0.8439, d = 0), suggesting a greater discrimination of cue and context stimuli.

### CCR5 amplifies anxiogenic behavior and diminishes exploration

To assess hyponeophagia, or the tendency to avoid risky behaviors such as eating in a novel environment, we used the novelty suppressed feeding (NSF) paradigm. Hyponeophagia is common in anxiogenic situations, including brightly lit and open spaces that rodents typically avoid due to perceived risk. The NSF is a conflict-based task in which mice are fasted overnight and then placed in a large, brightly lit arena for 10 minutes. Food is affixed to a central platform, and mice must linger in the center in order to eat. Compared to WT mice, which largely preferred to stay close to the arena wall for the duration of the test, CCR5 KO mice had a significantly lower latency to feed (*n* = 13 per group; *t*(24) = 2.702, *p* = 0.0125, *d* = 1.06) (Figure 1.B). In the elevated plus maze (EPM), a commonly used experimental paradigm that also relies on murine preference for dark, dimly lit places, CCR5 KO mice showed increased exploratory behavior, including a longer duration (*t*(20) = 2.217, *p* = 0.0384, *d* = 0.945) and increased mobility (*t*(20) = 2.162, *p* = 0.0429, *d* = 0.921) in the open arms (*n* = 11 per group; Figure 1.C). To ensure these changes were not due to strain differences in locomotor activity, we measured running wheel activity. For 9 days, mice were pair- or quad-housed in locomotion tracking cages (*n* = 12 per group). Running wheel activity was monitored using a beam-break sensor monitored by Scurry Activity Monitoring software to measure running duration and distance. Running wheel activity was averaged for each locomotion cage per genotype for each day. There were no significant differences in area under the curve (AUC) between genotypes (*t*(22) = 0.1902, *p* = 0.8509, *d* = 0.077) (Figure 1.D).

### CCR5 KO mice have stronger discrimination of cued and contextual stimuli

Previous studies have shown that CC5 homozygous and heterozygous mice show enhanced hippocampal contextual memory in a fear conditioning paradigm [13]. To support this data, and to assess cognitive flexibility in hippocampal learning, we also conducted fear conditioning, using a delay cued/contextual paradigm to extricate associative learning of a cued and contextual stimuli. During trial 1, mice were first exposed to a novel context (context A) for 6 minutes. During the first 3 minutes (stage 1), mice freely explored the chamber, to collect a baseline freeze response for each animal. Starting at minute 3, mice were administered a 30 second cue (70 dB tone), along with a co-terminating foot shock during the last 2 seconds of the cue. This repeated at the start of every minute, for a total of 3 times, to reinforce associated learning. After 12 days, mice were reassessed for both context and cue associations. For trial 2, mice were placed back into context A for 3 minutes with no foot shock or tone, and assessed for latency, duration, and episodes of freeze response. Following trial 2, mice were placed into a novel context (context B) for 3 minutes with the cue (70dB tone) to assess cue learning. To assess fear generalization, mice were allowed to explore context B for 1 minute after the cue ended. Results were z-score normalized to control for trial day variability for statistical analysis. Mixed model analysis (REML, Geisser-Greenhouse correction) showed a highly significant interaction between learning and genotype (*F*(3, 37) = 7.786, *p* = 0.0004) as well as strong affect of genotype alone (*F*(1, 24) = 20.89, *p* = 0.0001). Post-hoc analysis (Tukey’s HSD) showed that while genotypes did not differ at baseline during Trial 1 (*q*(23.02) = 0.1679, *p* = 0.9065, *d* = 0.031), CCR5 KO mice showed significantly higher freezing during both contextual (*q*(22.35) = 4.315, *p* = 0.0058, *d* = 1.137) and cue (*q*(4.625) = 4.168, *p* = 0.0352, *d* = 0.489) recall. To assess whether freezing was generalized to all novel contexts or cue specific, we compared freezing with and without the tone in context B, and saw no significant increase in generalization between groups(*q*(5.999) = 1.804, *p* = 0.2492, *d* = 0.825). When comparing tone versus no tone responses in Context B, CCR5 KO mice showed a greater discrepancy between tone and no-tone (*q*(7) = 2.331, *p* = 0.0526, *d* = 0.703) than WT (*q*(7) = 0.2044, *p* = 0.8439, *d* = 0) (supplemental figure 1A).

Overall, results did not imply increased generalization when compared to WT controls. CCR5 KO mice showed greater hedonic motivation, as well as neophilic exploration, when compared to WT mice in multiple distinct behavioral assays, suggesting behavioral translatability. Further, these results were not due to gross locomotor changes. As demonstrated here, as well as in previous studies, CCR5 KO mice exhibit stronger associative learning and greater discrimination of cue and contextual stimuli, suggesting stronger hippocampus-dependent memory and cognitive flexibility.

### CCR5 deletion increases dark cycle gamma power

Cortical network oscillations have long been studies for their role in information processing, mood regulation, and cognitive flexibility. We have previously shown that CCR5 knockout mice display enhanced gamma power and that CCR5 attenuation increases gamma oscillation power in mice harboring human apolipoprotein 4, a risk factor for Alzheimer’s disease [19]. To further explore the role of CCR5 in cortical network dynamics, we used *in vivo* intracranial wireless EEG telemeters with differential electrodes placed into the frontal and parietal lobes to record local field potentials from the cerebral cortex of freely behaving CCR5 KO and WT (*n* = 4 per group) mice (Figure 2.A). Continuous home cage recordings were collected for 5 days, and then segmented into dark and light cycles to analyze active period (dark cycle) oscillations. Due to the wealth of data supporting diminished gamma power in patients with MDD, as well as in murine models, we began by investigating broad cortical gamma power. Over the course of 5 dark cycles, CCR5 KO mice showed significantly higher gamma power (30-80Hz; *t*(6) = 2.867, *p* = 0.0285, *d* = 2.027) compared to WT mice. To confirm physiological relevance, we also compared light (inactive) cycle gamma power ((*t*(6) = 0.263, *p* = 0.8017, *d* = 0.186), and saw no significant differences (Figure 2.B). To further explore these gamma band differences, principle component analysis (PCA) was conducted on the mean dark cycle power spectral density (PSD). Surprisingly, PC1 explained 88.1% of the variance between groups (Figure 2.C). A closer look at PC1 indicated that the frequencies with the highest variance contributions were primarily in the gamma range (Figure 2.D). Spectral analysis of dark cycle recordings indicate that there were no significant power differences in any other low-frequency bands in the dark (delta, *t*(6) = 1.211, *p* = 0.271, *d* = 0.854; theta, *t*(6) = 2.056, *p* = 0.0855, *d* = 1.454; alpha, *t*(6) = 2.245, *p* = 0.0659, *d* = 1.588; *t*(6) = 2.251, *p* = 0.065, *d* = 1.592) (Figure 2.E) or light cycle (delta = *t*(6) = 0.305, *p* = 0.7706, *d* = 0.214; theta, *t*(6) = 0.221, *p* = 0.8325, *d* = 0.156; alpha, *t*(6) = 0.169, *p* = 0.8716, *d* = 0.119; beta, *t*(6) = 0.195, *p* = 0.8521, *d* = 0.138) (Supplemental figure 2A).

**Figure 2.**
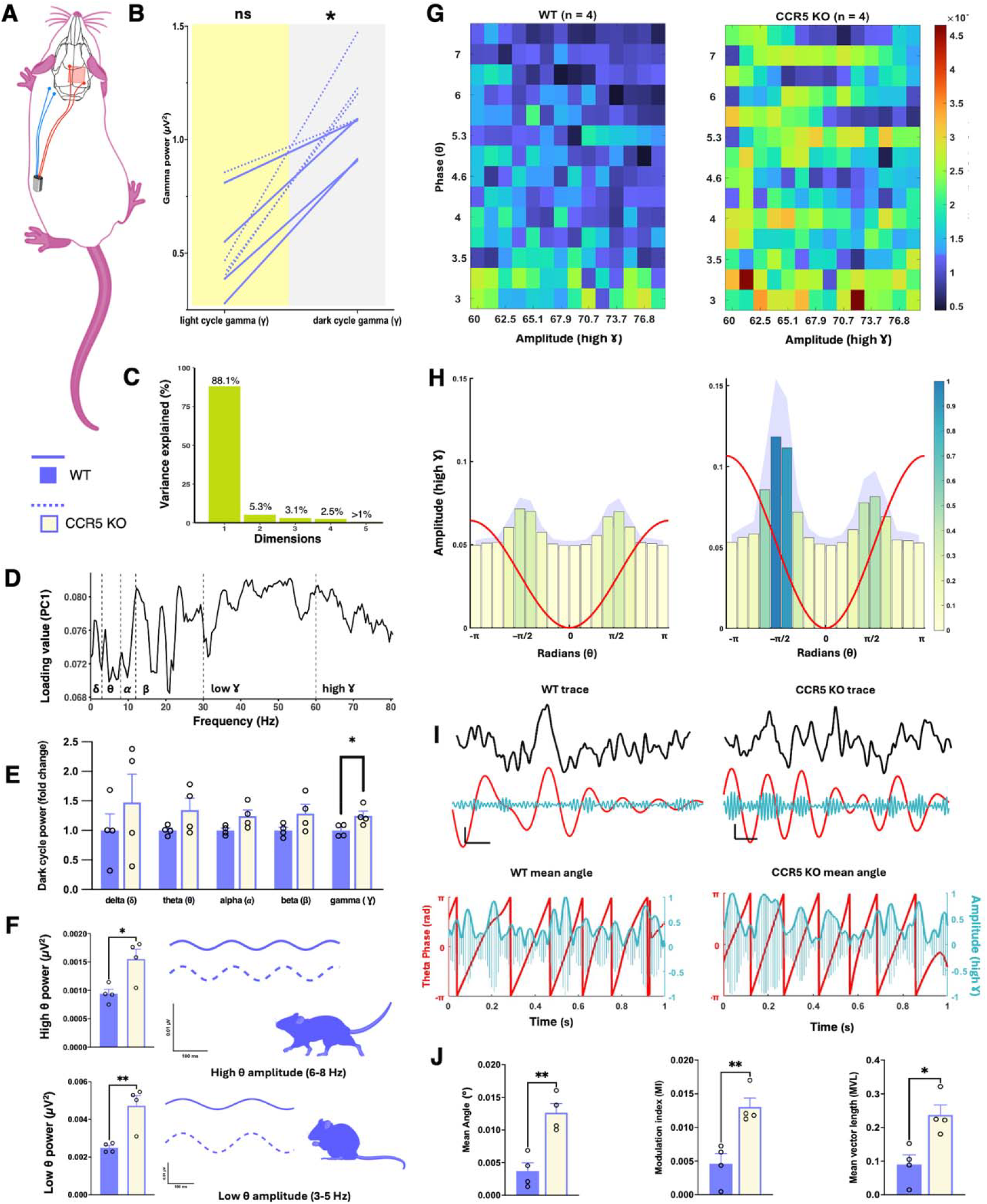
CCR5 constrains cortical oscillatory dynamics and cross-frequency coupling *in vivo*. Continuous intracranial EEG recordings for WT (*n* = 4) and CCR5 KO (*n* = 4) mice were conducted over 5 dark/light cycles using wireless epidural telemeters. **(A)** Schematic of telemeter and electrode placement. Epidural electrodes (DSI PhysioTel, HD-X02) were implanted in contralateral frontal and parietal regions. EMG leads were placed in the trapezius muscle, and the telemeter was implanted in the left flank. **(B)** Spectral analysis of gamma power showed significant changes in gamma power between WT (solid) and CCR5 KO (dashed) in the dark (active) cycle (*t*(6) = 2.867, *p* = 0.0285, *d* = 2.027), but not the light (inactive) cycle (*t*(6) = 0.263, *p* = 0.8017, *d* = 0.186). **(C)** PCA (correlation matrix, singular value decomposition) of dark cycle PSD (mV^2^/Hz; mean-centered, scaled to unit variance) attributed 81% of the variance between mean WT and CCR5 KO PSD to PC1. **(D)** Inspection of PC1 loading values showed that the largest contributors to PC1 were in the gamma band. **(E)** For statistical analysis, power was log-transformed and converted to a z-score to allow comparison between subjects. Note that graphs depict fold-change from control. Broad spectrum analysis indicated no significant changes in any other dark cycle (delta, t(6) = 1.211, p = 0.271, d = 0.854; theta, t(6) = 2.056, p = 0.0855, d = 1.454; alpha, t(6) = 2.245, p = 0.0659, d = 1.588; t(6) = 2.251, p = 0.065, d = 1.592) or light cycle (not pictured; delta = t(6) = 0.305, p = 0.7706, d = 0.214; theta, t(6) = 0.221, p = 0.8325, d = 0.156; alpha, t(6) = 0.169, p = 0.8716, d = 0.119; beta, t(6) = 0.195, p = 0.8521, d = 0.138) bands. Although there were no significant differences in broad spectrum theta, **(F)** burst analysis of slow (3-5 Hz) and fast (6-8 Hz) theta showed significant burst power differences (left) in both high/mobile (t(6) = 3.157, p = 0.0197, d = 2.233) and low/resting (t(6) = 3.966, p = 0.0074, d = 2.804) theta bursts. representative traces (right) show the amplitude differences between WT (solid) and CCR5 KO (dashed) theta bursts. To compare the phase-amplitude coupling (PAC) of theta and high gamma oscillations between groups, we calculated three measures of cross-frequency coupling: Modulation index (MI), mean vector length (MVL) and mean angle. **(G)** Heatmaps depict the mean PAC modulation index (MI) of WT (left) and CCR5 (right). **(H)** Phase-sorted amplitude shows the preferred theta phase (red line, radians) of the top 2% of high-frequency gamma oscillations. Darker phase bins indicate higher amplitude binning, and amplitude is indicated on the left y-axis. SEM is indicated by the purple shadow. **(I)** Representative traces (top; black) of WT (left) and CCR5 KO dark cycle recordings were transformed (middle; Hilbert) to extract the theta phase (red) and gamma amplitude (blue). The mean angle (bottom; red). **(J)** Statistical analysis of mean angle (t(6) = 4.736, p = 0.0032, d = 3.349), MI(t(6) = 4.19, p = 0.0058, d = 2.963), and MVL(t(6) = 3.552, p = 0.012, d = 2.511).Spectral bands = delta (0.5-2.5 Hz); theta (3-8 Hz); alpha (9-12 Hz); beta (13-29 Hz); gamma (30-80 Hz); low theta (3 – 5 Hz); high theta (6 – 8 Hz); low gamma (30 – 60 Hz); high gamma (60 – 80 Hz). All scale bars depict 100 μV by 100 ms.

### Burst, but not broad spectrum, theta power is increased in CCR5 knockout mice

While broad spectrum theta was not significantly different between WT and KO mice, the functional relevance of theta frequencies to MDD led us to conduct deeper analysis. Theta oscillations, particularly in the frontal and parietal cortex, can be locally generated or volume-conducted from the hippocampus. This phenomena leads to observable differences, both behaviorally and physiologically, between low (resting) theta, and high (mobile) theta. Further, these distinct theta oscillations are known to fire in complex burst patterns, and may desynchronize, which could diminish gross theta power [24]. To disentangle this, we analyzed both low and high theta for the first 12h dark cycle, which was chosen because the animals were relocated to a novel environment for recordings. We first identified potential bursts by filtering for either low (3-5 Hz) or high (5-8 Hz) theta using a 500 millisecond sliding window to identify short regions of substantially increased theta power. Bursts were categorized as theta events between 5 and 200 milliseconds, in which theta amplitude exceeded 3 standard deviations above the window mean. To ensure bursts were not over-counted, we included a minimum inter-burst interval of 200 milliseconds between the end of one burst, and the start of another. Finally, we filtered bursts against EMG activity to ensure physiological relevance. CCR5 KO mice showed significant differences in both high (*t*(6) = 3.157, *p* = 0.0197, *d* = 2.233) and low (*t*(6) = 3.966, *p* = 0.0074, *d* = 2.804) theta bursts, with the strongest differences in the low theta band (Figure 2.F).

### Theta gamma phase-amplitude coupling is weakened by CCR5

Finally, we analyzed cross-frequency phase-amplitude coupling of theta and gamma (PAC). PAC is particularly important to cognitive deficits observed with MDD, including reduced working memory [25]. The modulation index of theta-gamma PAC was significantly higher in KO mice across the full theta and full gamma spectrum (30-80 Hz) spectrum (Supplemental figure 2.B), but the strongest modulation occurred within the high gamma (60-80 Hz) band. We analyzed PAC strength using three metrics: Modulation index (MI), mean vector length (MVL), and mean angle. Group heat maps of WT and CCR5 KO mean MI show significantly higher modulation within the CCR5 KO group (Figure 2.G). The top 2% of gamma oscillations were binned into 20º bins to quantify the phase-sorted amplitude (Figure 2.H), and measure the MVL (Supplemental figure 2.C). To measure the mean angle, we extracted the theta and gamma oscillations from the EEG signal and calculated then phase angle differences (Figure 2.I). MI (*t*(6) = 4.19, *p* = 0.0058, *d* = 2.963), MVL (*t*(6) = 3.552, *p* = 0.012, *d* = 2.511), and mean angle (*t*(6) = 4.736, *p* = 0.0032, *d* = 3.349) were all significantly different between genotypes.

Taken together, these results suggest an activity-appropriate increase in gamma power, especially within high gamma frequencies, as well as burst-specific theta power.

### CCL5/CCR5 signaling inactivates GSK-3β and inhibits LTD

Previous studies have shown that CCR5 inhibition amplifies experience-dependent LTP in the barrel cortex through learning induced increases in mitogen activated protein kinase (MAPK) and c-AMP response element binding (CREB) protein activity [13]. Downstream targets of CCR5 include both G protein and β-arrestin pathways. To narrow our scope, we focused on arrestin signaling, because of its ability to attenuate GSK-3β [26], and subsequently inhibit LTD [27]. Importantly, LTD is thought to promote to cognitive flexibility, which is impaired in MDD [28].

To expand upon the role of the CCL5/CCR5 axis in GSK-3β signaling and LTD, we used primary hippocampal neurons from fetal rat or mouse cultures. First, we wanted to establish a timeline for CCL5/CCR5 activation of the glycogen-synthase kinase-3 beta (GSK-3β) pathway. Following CCL5 treatment (10 nM), neurons were collected, treated with a protease-phosphatase inhibitor, and lysed via rapid freeze-thaw to extract protein. Cells were collected in 20 minute intervals for 100 minutes, starting at 0 minutes. Western blot analysis (Figure 3.G) indicated a time-dependent increase in phospho-GSK-3β (pGSK-3β) which peaked at approximately 80 minutes, and was supported by a complementary increase in phospho-Akt/protein kinase B (PKB) (Figure 3.H).

**Figure 3.**
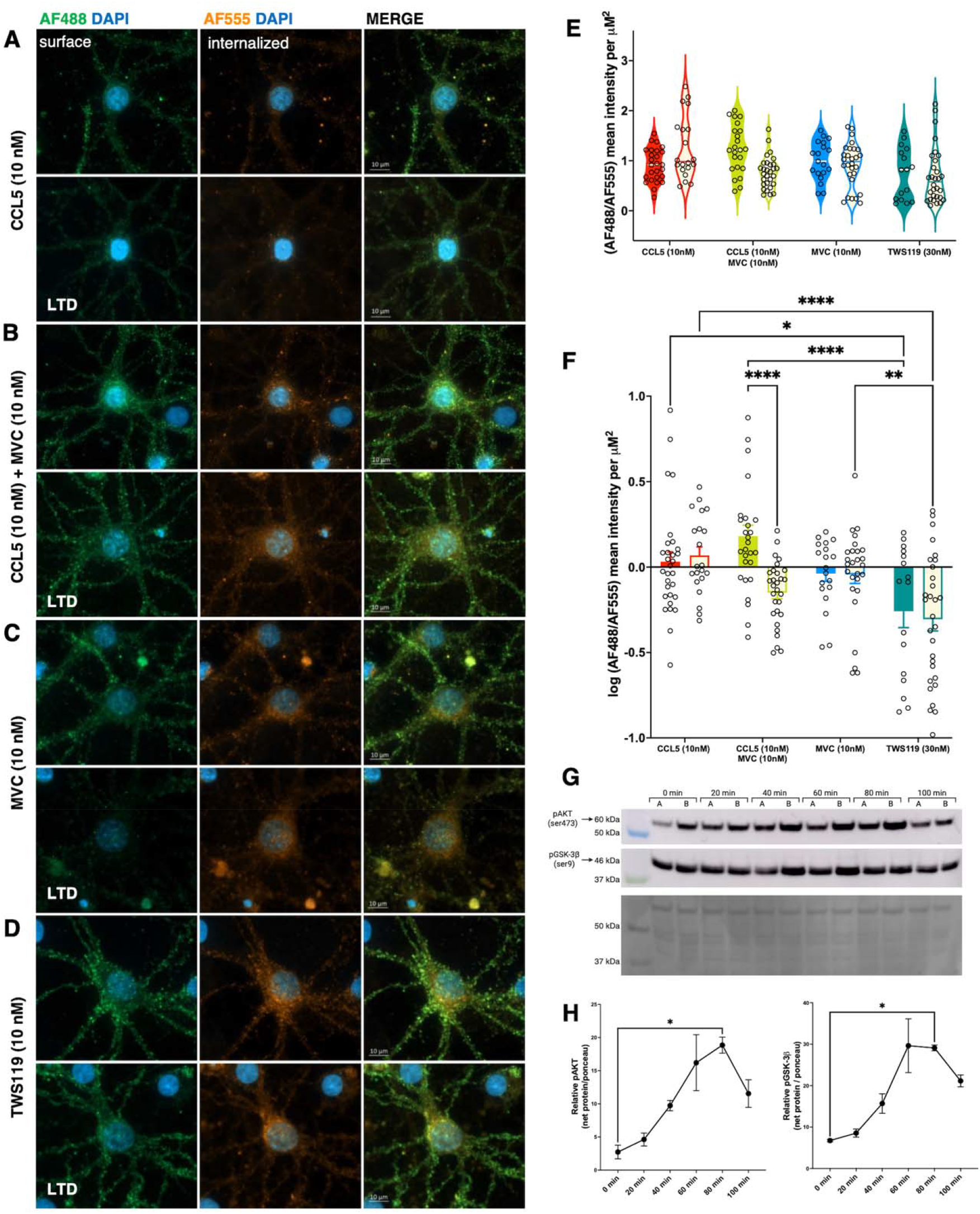
CCL5 impairs, and maraviroc rescues, LTD. Representative images showing immunofluorescent AMPAR internalization in primary rat hippocampal neurons. Neurons were pretreated with 10 nM of either CCL5 (A), CCL5+MVC (B), MVC alone (C), or a selective GSK3β inhibitor, TWS119 (D). For each treatment, LTD stimulated neurons (bottom pannel; 50 μM NMDA) were compared to treated, unstimulated controls (top panel). To quantify LTD, we labeled GluR2 (Millipore, MAB397) and quantified a ratio of surface (Alexa Fluor 488) AMPARs to internalized AMPARs (Alexa Fluor 555). The nucleus was visualized with DAPI. (E) Violin plots of non-transformed data show the distribution of mean intensity of external to internal staining in control and LTD-stimulated conditions (n=16-30 neurons per group, from 2 technical replicates per treatment and condition). (F) A 2-way ANOVA of Log(AF488/AF555) mean intensity per μM^2^ indicated significant internalization between control and LTD conditions in only the CCL5+MVC group. (*n =* neuron; 16-30 per group). (G)Western blotting DIV14 cortical neurons (top) normalized to total protein (ponceau, bottom) treated show a significant, time-dependent increase at 80 minutes post-CCL5 application (H).

The rapid inactivation of GSK-3β directed us to further investigate the role of the CCL5/CCR5 axis in synaptic plasticity. Long-term depression (LTD) is widely studied as a crucial counter-balance for long-term potentiation (LTP), and contributes to adaptive processes including cognitive flexibility [29], which is altered in MDD. To this end, we induced LTD in primary hippocampal rat neurons plated on glass coverslips using NMDA stimulation. Neurons were pre-treated with 10 nM of CCL5, maraviroc (MVC), a selective CCR5 antagonist, CCL5 and MVC (CCL5+MVC), or TWS119, a selective GSK-3β inhibitor known to induce AMPA receptor (AMPAR) internalization, which was used as a positive control. Surface AMPA receptors were labeled with a GluR2 primary monoclonal antibody, and LTD was induced using NMDA stimulation (50 μM). Surface and internalized AMPARs were differentially stained with AF488 or AF555 respectively, pre- and prior-to membrane permeabilization, to visualize LTD. A 2-way ANOVA of log-normalized mean intensity per μM^2^ revealed a highly significant main effect of both treatment (*F*(3, 192) = 11.31, *p* = < 0.0001, 13.69% variance) and LTD stimulation (*F*(1, 192) = 4.077, *p* = 0.0449, 5.142% variance). A highly significant interaction between group treatment and LTD condition (*F*(3, 192) = 4.249, *p* = 0.0062) accounted for 5.142% of the total variance (Figure 3.F). Post hoc analysis (Tukey’s HSD) of groups showed that AMPAR internalization, attributed to increased AF555 intensity, was not significantly changed in CCL5 (*q*(192) = 0.6242, *p* = 0.6594, *d* = 0.134), MVC-treated (*q*(192) = 0.09912, *p* = 0.9442, *d* = 0.02624), or TWS119 (*q*(192) = 0.7286, *p* = 0.607, *d* = 0.124) groups. It is notable that CCR5 antagonism restored receptor internalization in the CCL5 and MVC group (*q*(192) = 5.96, *p* = <0.0001, *d* = 1.209), highlighting the importance of both CCL5 and CCR5 in LTD (Figure 3.A – 3.D). Non-normalized mean intensity shows the distribution of internalization ratios within groups (Figure 3.E).

Although it is known that CCL5 can bind to a variety of receptors including CCR3 and CCR1 [30], LTD was rescued by MVC, a highly selective CCR5 antagonist. These results suggest that CCL5 impairs AMPAR-mediated LTD through a CCR5-dependent GSK3-β pathway.

### CCR5 bi-directionally modulates PV expression

Though prior work has examined the effects of CCL5 on excitatory neurons, its effects on parvalbumin expressing inhibitory neurons has not been well-examined. GABAergic interneurons, especially parvalbumin positive (PV+) interneurons, modulate excitatory firing of PCs and play a key role in the generation of gamma oscillations. Due to the direct role of PV+ interneurons in excitatory / inhibitory (E/I) maintenance and gamma oscillations, we sought to understand whether CCR5/CCL5 signaling plays a role in PV+ expression. To establish whether CCR5/CCL5 could have a direct effect parvalbumin expression, we co-stained primary hippocampal TCs for PV (AF555) and CCR5 (AF647), along with MAP2 (AF488) and DAPI to identify neurons. We observed CCR5 expression on both PV+ and PV-neurons, and interestingly, found the intensity to be highest on the soma of both neuron types (Figure 4.A). To look at the effects of CCL5/CCR5 signaling, we again treated neuronal cultures with CCL5 (Figure 4.C), CCL5 and MVC (Figure 4.D), and DMSO (supplemental figure 4.A). A 2-way ANOVA (*n* = 10 neurons per group, per timepoint), showed a significant main effect of CCL5 treatment (*F*(2, 108) = 13.95, *p* = <0.0001, 18.87% variance explained). Following CCL5 treatment, PV+ neurons had a time-dependent increase in PV intensity, with expression increasing most significantly between 0 and 24 hours (*q*(108) = 5.27, *p* = 0.0009, *d* = 1.4827) compared to CCL5 + MVC neurons (Figure 4.B). Western blot analysis of primary hippocampal neurons at 0 and 24 hours showed the same effect (*q*(12) = 3.065, *p* = 0.051, *d* = 2.685) (Supplemental figure 4.B). These results are consistent with the fact that phosphorylation of GSK-3β at serine 9 deactivates GSK activity, as well as with published work showing that PV-specific deletion of GSK-3β increases PV activity [31].

**Figure 4.**
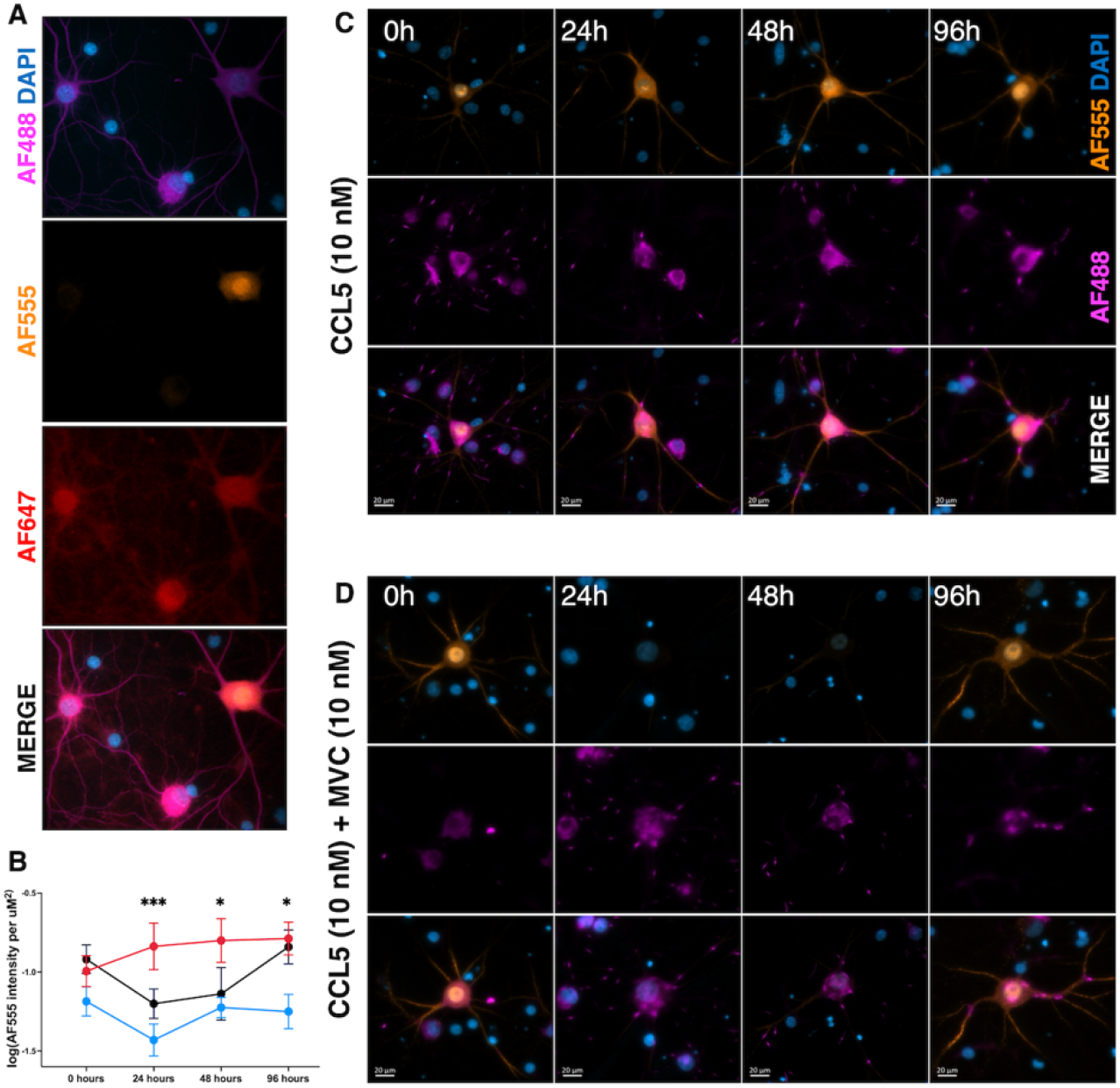
CCL5 and maraviroc have oppositional effects on parvalbumin expression. **(A)** CCR5 (AF647) is expressed on both PV+ (AF555) and PV-neurons (MAP2 = AF488) in primary hippocampal cultures. **(B)** Parvalbumin is bidirectionally regulated by CCR5/CCL5 activation in a time-dependent manner. In a 2-way ANOVA, CCL5 treatment showed a highly significant main effect (F(2, 108) = 13.95, p = <0.0001, 18.87% variance explained). CCL5 treated neurons (red) showed significantly higher parvalbumin expression compared to, parvalbumin expression was upregulated at 24 (q (108) = 5.27, p = 0.0009, d = 1.4827), 48(q(108) = 3.773, p = 0.0237, d = 1.249), and 96 (q(108) = 4.108, p = 0.0123, d = 1.3741), but not 0 (q(108) = 1.701, p = 0.4539, d = 0.635) hours. Intensity was measured on 10 neurons per group, per time period. Representative images of CCL5 and CCL5 + MVC treated images **(C-D)** show parvalbumin expression (AF555) at 0, 24, 48, and 96 hours.

### CCR5 KO mice show reduced hippocampal *Gabra4* and tenascin C, two molecules that can inhibit pyramidal cell excitability

To understand how CCR5 deletion alters the hippocampal proteome, we conducted an unbiased proteomic analysis of CCR5 KO and WT mice on hippocampal lysates, which were collected approximately 4 hours after fear conditioning recall. Results of data independent acquisition (DIA) gave a total of 4124 detected proteins. We leveraged two separate statistical methodologies to categorize significant differences in our dataset. First, we used Spectronaut (Biognosys) software to quantify significant protein changes between CCR5 and KO mice using an unbiased approach. Then, using a training dataset comprised of fear conditioning data, as well as immunostaining of E/I proteins of interest, we used a logistic Bayesian model to incorporate a robust and targeted exploration of cognitive associated proteins. STRING database analysis (Version 12.0) of significantly changed proteins showed significant enrichment in all datasets (Spectronaut, *n* = 713 proteins, p_enrichment_ < 31.0 e^−16^; Bayesian, *n* = 309 proteins, p_enrichment_ = 6.63 e^−6^; Overlap, *n* = 231 proteins, p_enrichment_ = 3.59 e^−6^) (Figure 5.A-B). Of interest were tenascin C and GABRA4, two proteins that play a role in maintenance of inhibitory synapses (Figure 5.C). Interestingly, levels of tenascin C, a component of perineuronal nets which generally enhance PV activity [32], was significantly downregulated in CCR5 KO mice. Notably, both chondroitinase mediated PNN digestion and specific knock out of tenascin C have been shown to increase gamma power in murine models [33, 34]. Further, reductions in extracellular matrix can also enhance cognitive flexibility [35]. The GABA-A α4 (GABRA4) receptor, which is localized to the extrasynaptic space, was also significantly reduced in CCR5 KOs. This GABA receptor subunit is expressed on pyramidal cells and plays a prominent role in tonic inhibition of pyramidal cells. Increases in tonic inhibition have also been linked to reduced PC intrinsic excitability and LTP [36, 37].

**Figure 5.**
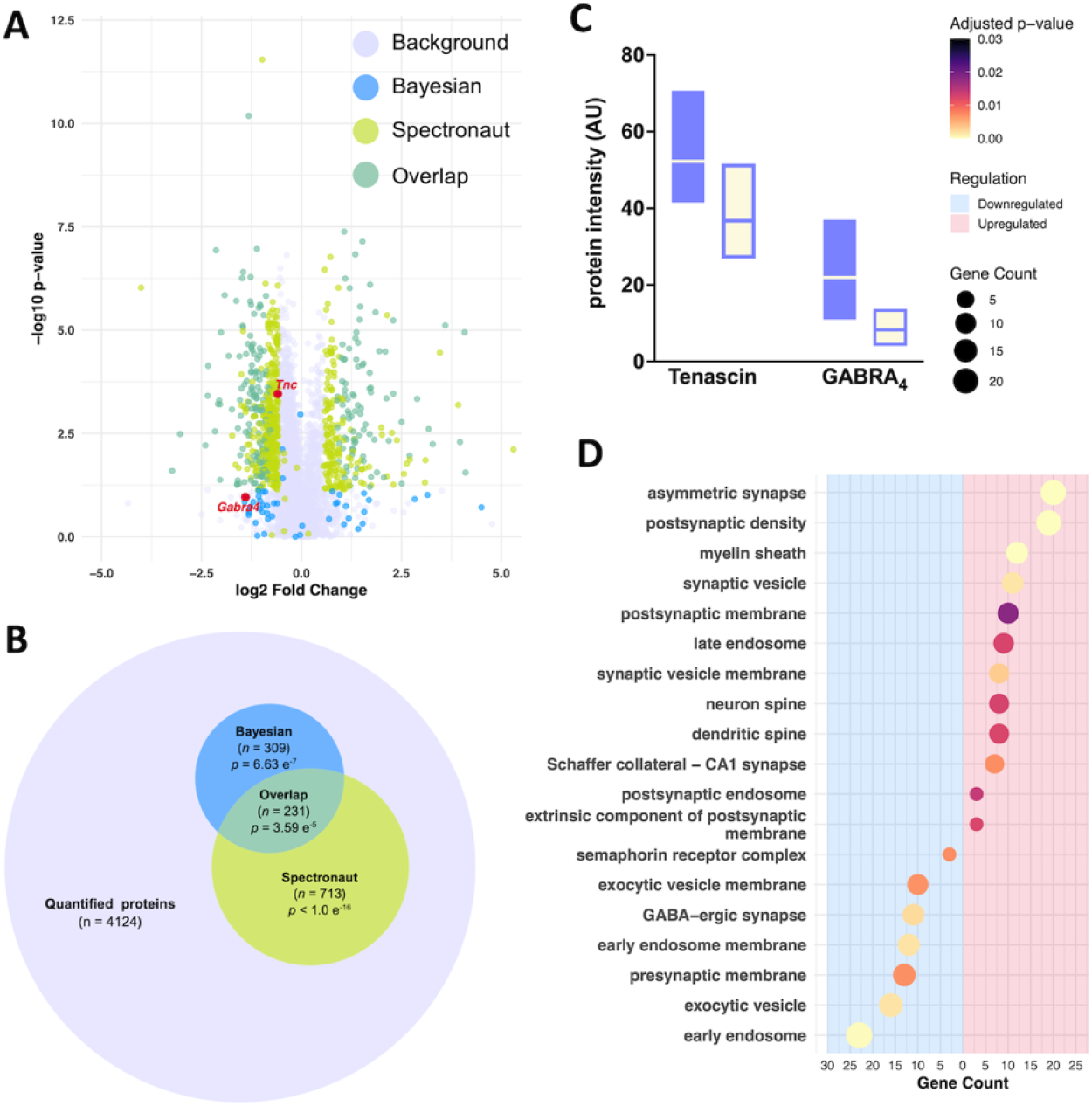
CCR5 KO mice show substantial differences in plasticity-associated protein expression after a learning event. Proteomic results indicate better hippocampal synaptic plasticity. (A) Volcano plot of proteomic results of hippocampal lysates (*n* = 5 per group) show the results of statistical analysis. (B) A Euler plot depicts relative size and overlap of significant findings. A total of 4124 proteins were quantified (light purple). Results from Spectronaut DAI (green) showed 713 significantly changed proteins (STRING PPI enrichment p-value < 31.0 e^−16^). Bayesian analysis was completed using a training dataset generated acquired from delay cued/contextual fear conditioning, as well as hippocampal immunostaining data (Supplemental Table 1), and showed an additional 309 significantly changed proteins (purple, STRING PPI enrichment p-value = 6.63 e^−6^). 231 proteins overlapped between analysis (teal, STRING PPI enrichment p-value = 3.59 e^−6^). (C) GABRA4 (*p*_*bayes*_ *=* 0.0141, *q*_*bayes*_ *=* 0.23047981, *p*_*Spectronaut*_*=* 0.11016626, *q*_*Spectronaut*_= 0.06909672, −62.17% change in CCR5 KO mice) and tenascin (*p*_*bayes*_ *=* 0.0241, *q*_*bayes*_ *=* 0.2604,*p*_*Spectronaut*_*=* 0.0003, *q*_*Spectronaut*_= 0.0013, −33.69% change in CCR5 KO mice), two proteins of interest, were both significantly downregulated in CCR5 KO mice. (D) GO enrichment analysis of cellular components showed a pattern of up- and down-regulation that indicates increased neural branching, asymmetric (neuron-neuron) synapses (Fold-enrichment = 4.913, *z* = 7.995, adjusted *p* = 1.210e^−6^), and post-synaptic density (Fold-enrichment = 4.908, *z* = 7.785, adjusted *p* = 1.639e^−6^)as well as CA1 synapses(Fold-enrichment = 3.964, *z* = 4.993, adjusted *p* = 0.007). GABAergic synapses(Fold-enrichment =5.545, *z* = 5.141, adjusted *p* =0.002), pre-synaptic membrane (Fold-enrichment = 2.941, *z* = 4.134, adjusted *p* = 0.006), and exocytic vesicles (Fold-enrichment = 3.210, *z* =5.001, adjusted *p* =0.001), were downregulated. A full list of enrichment scores are available in Supplemental Table 2.

**Figure 6.**
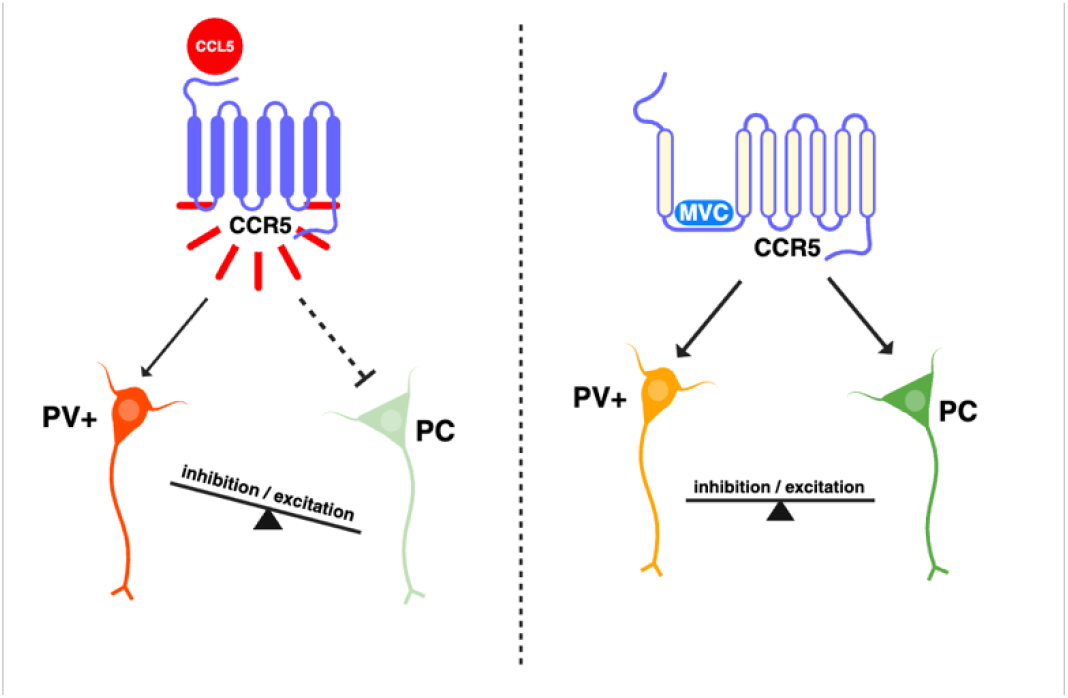
CCL5 and CCR5 bidirectionally modulate E/I balance. CCL5/CCR5 agonism directly upregulates parvalbumin levels on GABAergic PV+ interneurons, increasing the inhibitory modulation of PV+ interneurons on excitatory PC cells. Inhibiting CCR5 with maraviroc restores E/I balance by normalizing parvalbumin and regulating inhibitory control of PCs.

### CCR5 deletion shifts hippocampal synapses towards excitability and potentiation

To look at large-scale alterations in signaling and plasticity, we conducted GO enrichment analysis of cellular components. CCR5 KO mice showed significant bi-directional enrichment in pro-plasticity cellular components (Figure 5.D). Interestingly, CCR5 KO mice showed significant connectivity differences, including increased neuron-to-neuron synapses (asymmetrical synapse fold-enrichment = 4.913, *z* = 7.995, adjusted *p* = 1.210e^−6^), especially in proteins specific to Schaffer’s collateral CA1 synapses (Fold-enrichment = 3.964, *z* = 4.993, adjusted *p* = 0.007), which are primarily glutamatergic and play a crucial role in LTP. Concurrent downregulation of GABAergic synapses (fold-enrichment = −3.965, *z* = 4.994, adjusted *p* = 0.002) supports a general increase in hippocampal and cortical excitability. In addition, upregulated neuronal processes (neuron spine fold-enrichment = 3.858, *z* = 4.154, adjusted *p* = 0.012; dendrite spine fold-enrichment = 3.938, *z* = 4.223, adjusted *p* = 0.012) and extrinsic components of the post-synaptic membrane (fold-enrichment = 14.953, *z* = 6.278, adjusted *p* = 0.012), suggests increased synaptic tone. Downregulation of the semphorin 4D receptor complex (fold-enrichment = −16.908, *z* = 6.756, adjusted *p* = 0.007), which promotes inhibitory synapse formation, reinforces a shift away from GABAergic signaling. Of interest, post-synaptic upregulation (postsynaptic density fold-enrichment = 4.908, *z* = 7.785, adjusted *p* = 1.639e^−6^; postsynaptic membrane fold-enrichment = 2.946, *z* = 3.626, adjusted *p* = 0.018) was also observed, as was an increase in synaptic vesicle proteins (synaptic vesicle fold-enrichment =4.553, *z* = 5.573, adjusted *p* = 0.001; synaptic vesicle membrane fold-enrichment = 5.563, *z* = 5.512, adjusted *p* =0.003).

In the context of memory and cognitive flexibility, these proteomic results suggest a highly plastic hippocampal environment in CCR5 KO mice that is suited for rapid, dynamic modulation following a hippocampal-dependent learning event.

## Discussion

Studies have shown that CCL5 is elevated in patients with MDD, as well as in conditions that can increase MDD risk, including chronic HIV infection and Alzheimer’s disease [10–12, 38–40]. Evidence suggests that chronic cytokine upregulation is associated with HPA axis sensitization, as well as pervasive symptoms and treatment resistance, which supports the investigation of anti-inflammatory drugs for antidepressant effects [41]. To better inform the question of whether CCR5 antagonists should be considered as an adjunct therapy for specific populations, the present study focused on the possibility that CCL5 signaling through the CCR5 receptor might contribute, in some part, to MDD-associated symptoms and treatment resistance.

Behavioral studies comparing strain matched wild type and CCR5 knockout mice suggested that CCR5 presence increased behavioral correlates of depression including anxiety, anhedonia, and impaired cognitive flexibility. Select behavioral effects are consistent with published work in other preclinical and clinical studies, including work showing that CCR5 deletion in mice improves LTP and the duration of memory linking [42], as well as work in humans suggesting that maraviroc can improve ratings on the Montgomery Asberg depression rating scale in patients with post-stroke depression [43]. This study provides preclinical evidence to demonstrate that CCR5 signaling may also elevate levels of anxiety and anhedonia, specific endpoints that can further impact quality of life in patients with MDD.

In addition to behavioral alterations, an increasing body of evidence suggests that MDD is also associated with EEG changes that may underlie select behavioral and cognitive alterations. In particular, studies conducted in both rodent models and humans suggest that the power of cortical high frequency gamma oscillations may be reduced in MDD, and improved with remission of symptoms and/or effective antidepressant treatment [5, 14–16, 20]. Moreover, in a stress model of murine depression, *ex vivo* hippocampal gamma power was reduced [5].

Previously, we reported that CCR5 attenuation improved gamma power in apolipoprotein 4 (Apo4) targeted replacement mice. We observed increased 20-55 Hz gamma power in CCR5 knockouts when compared to wild type controls, and also observed that CCR5/ApoE3 heterozygotes had increase gamma power in this range when compared to CCR5/ApoE4 heterozygotes [19]. In the present study we examined a second cohort of wild type and CCR5 knockouts for gamma power in both the low gamma power range and in the 60-80 Hz range. We also extended EEG studies to examine theta gamma phase amplitude coupling (PAC), which is altered in MDD and potentially important to information transfer [44]. EEG studies suggested that CCR5 attenuation increases gamma power across a broad range of frequencies but more significantly in the 60-80 Hz range, and that CCR5 attenuation also increases PAC and theta burst power. A general increase in cortical E/I balance could contribute to increases in both gamma power and PAC, and both have the potential to enhance cognitive processing [45, 46].

Biochemical studies described herein further demonstrated that CCL5 stimulation increased phosphorylation of GSK-3β at serine 9, a site at which phosphorylation has been shown to inhibit GSK-3β activity [47]. Though we did not address the underlying mechanism, based on prior studies a likely candidate is CCR5-coupled b-arrestin signaling [48]. The timing of GSK-3β phosphorylation in our studies is also consistent with β-arrestin signaling. Importantly, in studies from the Collingridge lab, GSK-3β activity has been shown to be critical to NMDA dependent LTD, an important form of plasticity for cognitive flexibility that is also impaired in MDD [49]. We thus collaborated with the Pak lab at Georgetown, which regularly examines NMDA-dependent LTD, to examine CCL5’s effect on this form of plasticity. Experimental results suggested that CCL5 impaired LTD in a manner that was blocked by maraviroc and thus CCR5 dependent. Despite the implications for potential adverse effects of LTD increases with CCR5 antagonists, however, the finding that fear memory linking is increased by attenuation of CCR5 signaling suggests that adaptive memories may be spared [42].

Our cell culture based studies also demonstrated that CCL5 stimulation of cultured neurons increased PV expression, as detected by immunostaining and Western blot analysis. The time course of this increase was consistent with prior work examining PV changes in response to learning [50]. While PV levels in the CCR5 knockouts were on average lower than those is the wild type with proteomics analysis, the difference was not significantly different. Non-significant changes in proteomics could, however, reflect a relatively low sample number as well as the possibility that endogenous CCR5 agonist levels low or varied in the wild type mice. Published work suggests that increased PV expression, and aggrecan levels in particular, is generally correlated with increased PV neuronal activity, and thus expression is often used as one proxy of the same [51–55].

Potential changes in PV expression and PV neuronal activity are of interest. While previous work has demonstrated that CCL5/CCR5 axis signaling can suppress PC activity, its potential to increase PV activity may also contribute to CCL5’s effect on overall E/I balance. Future studies should thus examine CCL5’s effects on PV activity more directly through patch clamp and/or targeting calcium imaging studies. It is tempting to speculate that gamma changes and the ability of CCR5 antagonists or the functional human CCR5 deletion (CCR5 delta 32 variant) to improve post stroke outcome could follow from increased pyramidal plasticity that occurs with a generalized increase in cortical E/I balance [43]. It is also of interest to hypothesize that the potential for CCR5 antagonism to disinhibit pyramidal excitation could represent a mechanism of antidepressant efficacy that is shared with first line antidepressants including fluoxetine and venlafaxine [5, 56].

Proteomics changes showed significant differences between the ECM/PNN component tenascin C levels in wild type and CCR5 knockouts, with reduced tenascin C in the knockouts. This is consistent with the previously described ability of CCR5 antagonists to reduce ECM deposition in varied end-organs [7, 8], as well as to stress/inflammation based changes in ECM [9, 57]. This is also of interest to potential effects of CCR5 on inflammation and PV interneuron activity in that tenascin C can increase the former [58], and tenascin C may also contribute to stability of the PNNs that impact PV activity [59, 60]. Of note, the first line antidepressant fluoxetine also acts in part through its ability to reduce PNN levels and promote reversal learning [17].

Of additional interest, a Bayesian-based proteomic analysis also showed that Gabra4 levels were reduced in CCR5 knockouts. Gabra4 is extra-synaptic in localization and contributes to tonic inhibition of the pyramidal cells on which it is expressed. Increases in tonic inhibition have also been linked to reduced PC intrinsic excitability and LTP [36, 37], and thus it is tempting to speculate that reduced Gabra4 levels in the knockouts could contribute in part to the previously described memory skills in CCR5 KO animals [13].

Overall the data herein suggests that CCL5/CCR5 signaling affects behavioral and neurophysiological changes, including increased theta-gamma phase amplitude coupling, that are relevant to MDD. CCL5/CCR5 signaling may further affect both LTP and LTD, two important forms of plasticity that can be impaired in MDD. CCL5/CCR5 axis-dependent effects on GSK-3β activity and LTD, as well as potential effects on tonic inhibition, may in turn contribute to the cognitive inflexibility and perseveration that can affect select individuals with MDD [61, 62]. Accordingly, available oral CCR5 antagonists could be increasingly considered for post-stroke depression, HIV-1 infected patients with depression, and as potential adjunct therapy in select patients with MDD.

## Materials and Methods

### Animals & Behavioral assays

Mice were adult (4-6 months) male CCR5 knockout (CCR5 KO, JAX #005427) or wildtype (WT, JAX #000664) C57BL6J mice from The Jackson Laboratory. Animals group-housed with enrichment (hiding structure, nestlet) in a typical 12 hour light/dark cycle with food and water *ad libidum* unless otherwise specified. Animals were not subjected to additional stressors, in that traditional housing is associated with stressors including limited space, poor enrichment, potential territorial confrontations, and tail handling during cage changes. Moreover, prior studies identifying biochemical mechanisms and depression related behavioral effects of fluoxetine and other monoamine modulating antidepressants did not include additional stressors [17, 18].

### Behavioral assays

Behaviors were conducted in experimentally naive animals during the light cycle. For all behavioral tasks, mice were habituated to handling, as well as the test room, prior to the start of the task. Anxiety assessments, including novelty suppressed feeding (NSF) and the elevated plus maze (EPM), were conducted in a brightly lit room to increase the contrast between shadowed and illuminated regions of the test apparatus. Fear conditioning was conducted in sound and light-proof chambers. The sweet drive test (SDT) was conducted during the dark/active cycle in red light. For the NSF and SDT, mice were fasted for approximately 16 hours before the test.

#### Locomotion

Mice were pair- or quad-housed in locomotor cages with beam-break running wheels equipped with Scurry Activity Monitoring Software (Lafayette Instruments, Lafayette, IN, USA) for 10 days in a typical 12-hour light/dark cycle to measure locomotor activity.

#### EPM

Mice were placed in the center zone of the maze and allowed to explore for 5 minutes. Exploration was tracked using ANYMaze (version 7.2). Zone entries, distance, and time in zone were tracked using the center or the animal.

#### NSF

Mice were placed into the corner of a large (24×36 inch) brightly lit arena with a chow pellet affixed to a central platform. Test duration was 10 minutes. Latency was scored separately by two researchers, and averaged. Feeding latency was measured as the interval from the trial start to the time when the mouse approached the central platform and took a bite of the chow pellet. Movement was tracked using ANYMaze.

#### Fear conditioning

Mice were habituated in habituation room A (dim white light, white noise, lemon scent) before being placed into the fear conditioning chamber in context A (lemon scent, white walls, palm handled, slatted floor) for trial 1. For trial 1 stage 1, mice were allowed to explore the chamber for 3 minutes. For trial 1 stage 2, a 30 second tone (cue) was paired with a mild foot shock (0.3 mA) that was administered during the last 2 seconds of the tone, followed by a 30 second interval with no cue or foot shock. This was repeated for a total of 3 times. After 12 days, all mice were placed back into Habituation room A before completing trial 2. For trial 2, mice were placed back into context A for 3 minutes, with no cue or foot shock. After trial 2, mice were immediately returned to their home cages and moved to habituation room B (mint scent, no noise, no light) for 90 minutes. Then, they were placed into the fear chamber in context B (mint scent, checkered walls, solid floor, tunneled into chamber), and recorded for 3 minutes while the cue (tone) was played. Finally, for 1 minute, mice explored context B with no tone. Freezing time, latency, and episodes were tracked using ANYMaze.

#### SDT

The sweet drive test is a modified sucrose preference test that uses a food reward. For 2 days, mice were habituated to a sweet grain-based cereal as well as the chamber apparatus, which was a modified T-maze with no internal gates/walls. For the test, mice were placed in the starting arm of the T-maze, and allowed to explore freely for 10 minutes. In one arm, a pre-weighed 500 mg chow pellet was placed. In the other, 500 mg of cereal was placed. Food placement was switched after each trial to mitigate location preference. Food consumption and first side of entry were measured. Animals completed 3 trials, 48 hours apart.

### Electroencephalography (EEG) Recordings

#### Telemetry implantation surgery

Prior to implantation, animals were group-housed and acclimated to handling. Wireless implantable DSI PhysioTel HD-X02 telemeters (DSI, St. Paul, MN, USA) were used to record cortical activity. Stereotaxic surgery was performed by a veterinary surgeon using a well-established surgery protocol [19, 20]. Briefly, mice were anesthetized with continuous inhaled isoflurane and secured in a stereotaxic device with a heating pad. Transmitters were placed subcutaneously near the left hind flank. Two electromyography (EMG) leads were placed in the cervical trapezius muscle to capture locomotor activity. Two intracranial EEG leads were secured with epidural screws, placed contralaterally in the frontal (1mm anterior, 1mm lateral) and parietal (3mm posterior, 1mm lateral) lobes. Electrodes were secured and isolated from tissue with dental cement. Analgesics (carprofen) was administered pre- and post-operatively for pain relief. Prophylactic antibiotics (Baytril, subcutaneous injection; triple antibiotic ointment, topical) were administered for 3 days via injection following surgery. Sutures were removed after 7 days. Animals recovered for 1 week before recordings.

#### Signal acquisition & preprocessing

For recordings, mice were moved to a novel room and single-housed with typical conditions in clear cages placed on top of a wireless DSI PhysioTel RPC-1 receiver connected to an MX2 matrix (sampling rate = 500 Hz). Electrode location was confirmed again post-euthanasia, to ensure accurate placement. LabChart was used to record signals continuously for 5 dark/light cycles. Signals were preprocessed using LabChart (AD Instruments, Colorado Spring, CO, USA) to filter signals for the DSI-validated accuracy range (0.5 – 80 Hz). Artifacts were removed using the EEGLAB Toolbox (v2024.1) in MATLAB (R2024a).

#### Power analysis & power spectral density (PSD)

Absolute power was were calculated for each 12-hour dark and light cycle using LabChart 8 Reader. Power spectral density (PSD) was calculated in MATLAB.

#### Theta-gamma phase-amplitude coupling (PAC)

PAC was calculated for the first 12-hour dark cycle using the PACT toolbox in EEGLAB. Signals were filtered for theta and gamma frequencies. A threshold of 2% of highest amplitudes was used to obtain the HAS index.

Modulation index (MI), mean vector length, and mean angle were calculated using a total of 500 surrogations (statistical control), and phase was binned into n=18 bins.

#### Theta burst analysis

To isolate theta burst events, the signal was band-pass filtered at either high or low theta, and the analytic signal (Hilbert) was calculated. A 500 millisecond sliding window was used to set a dynamic threshold of three standard deviations above the mean envelope amplitude, to identify segments of potential burst firing. Criteria for bursts were a duration between 5 and 200 milliseconds, as well as a minimum inter-burst interval of 200 milliseconds, and an EMG filter (low theta = immobile; high theta = mobile) to ensure physiological relevance. Mean frequency and power were calculated for all burst events. Principle component analysis (PCA) for PSD was conducted in R.

### Tissue preparation

Mice were deeply anesthetized using inhaled isoflurane, followed by a toe pinch to confirm non-responsiveness, and then rapid decapitation. All tissue prep was completed on ice. Brains were extracted and hemisected along the sagittal plane.

#### Hippocampal lysate preparation

For protein extraction, the hippocampus was sub-dissected and sonicated in 200 μL lysis buffer (RIPA cell lysis buffer [50 mM Tris, pH 7.5, 150 mM NaCl, 0.1% sodium dodecyl sulfate, 1% octylphenoxypoly (ethyleneoxy) ethanol, branched], 1x protease & phosphatase cocktail [Halt, Ref#1861281]) for 30 seconds in 3 second pulses before being centrifuged at 13.3x g for 10 minutes at 4°C. Supernatants were removed, flash-frozen in dry ice, and stored at −80°C until use. Total protein was determined using the Pierce BCA Protein Assay Kit (Thermo Scientific, Cat#23225) following the microplate procedure.

### Primary hippocampal cultures

#### Culture preparation

Tissue cultures (TCs) were prepared according to previously published methodology used in the Pak lab [21]. In short, pregnant Sprague-Dawley rats (8-10 week dams) from Charles River (Raleigh, NC, USA) were euthanized at embryonic day 18 (E18) using flow-regulated inhaled carbon dioxide, followed by toe pinch and cervical dislocation. Hippocampal neurons from E18 rat embryos were collected (approximately 8-14 embryonic hippocampi per culture) and plated on poly-D-lysine/lamanin coated coverslips (Sigma, cat. #P8099, Sigma, cat. #L2020) in a 12-well plate at a density of 75,000 cells/well. Cultures were grown in supplemented neurobasal media (Gibco, cat# 21-103-049, [SM1 (STEMCELL Technologies), primocin (InvivoGen, #ant-pm-05), 0.5□mM glutamine, 12.5□mM glutamate]). Media was changed at day in vitro (DIV) 5 and 12. Experiments were conducted between DIV 21-28. All drugs and antibodies, and timepoints for tissue culture experiments are detailed in Supplemental Table 1, unless otherwise specified.

#### CCL5 time course experiments

All drugs were diluted in DMSO. Neurons were treated with CCL5 (10 nM), maraviroc (10 nM), CCL5 and MVC (10 nM each), or DMSO alone. After pretreatments were applied, coverslips or western blot lysates were collected in intervals starting at T0 (0 minutes).

#### LTD stimulation

Wells were pre-treated for 45 minutes with CCL5 (10 nM), MVC (10 nM), CCL5 and MVC (10 nM each), or TWS119 (30 nM). Then, coverslips were incubated with a mouse anti-GluR2 (10 μg/mL) at 37°C for 10 minutes to label AMPA receptors. After briefly washing with DMEM, LTD was stimulated by incubating coverslips in NMDA (50 μM) for 5 minutes before being placed back in pre-treated conditioned medium for 25 minutes to allow for receptor internalization. Control groups were not LTD-stimulated.

#### Immunocytochemistry

For immunocytochemistry (ICC), coverslips were rapidly placed into fixation medium (4% paraformaldehyde, 4% sucrose in 1xPBS) for 5 minutes and washed 3x for 10 minutes in PBS. Neurons were incubated for 10 minutes with a permeabilizing background suppressor (Biotum TrueBlack IF Background Suppressor System (permeabilizing), Cat# 23012), before being washed 3x for 10 minutes in PBS and blocked for 1 hour at room temperature in blocking buffer (1x PBS + 3% normal goat serum + 0.1% BSA) and incubated overnight in the appropriate primary antibody in a humidity chamber at 4°C. The next morning, coverslips were washed 3x for 10 minutes in PBS and incubated with the appropriate secondary for 2 hours at room temperature in a humidity chamber. For Internalization staining, coverslips were rapidly placed into fixation medium (4% paraformaldehyde, 4% sucrose in 1xPBS) for 5 minutes and washed 3x for 3 minutes each in PBS. Secondary fluorophores were diluted in 3% normal goat serum (Invitrogen, cat#50062Z), 0.1% BSA in 1x PBS. Surface AMPARs were labeled with AF488 goat anti-mouse (1:200) by incubating coverslips for 2 hours at room temperature. After surface labeling, the membrane was permeabilized by rapidly submerging coverslips in −20°C methanol for 90 seconds, and washed 3x 5 minutes in 1xPBS. Internalized AMPARs were labeled with AF555 goat-anti-mouse (1:300) by incubating coverslips for 1 hour at room temperature. All coverslips were washed 3x 10 minutes and briefly rinsed with distilled water before mounting with mounting media with DAPI (Vectashield, H-1500).

#### TC Lysate preparation

For western blot lysates, media was rapidly aspirated with vacuum suction and replaced with 200 μL of lysis buffer. Wells were thoroughly scraped with a sterile cell scraper and lysis buffer was gently pipetted on walls and swirled to collect all cells before lysates were placed on ice. Once all lysates were collected, tubes were inverted using a tube rotator for 10 minutes at 4°C to further lyse cells. Proteins were extracted via rapid freeze-thaw (2 minutes on dry ice followed by 2 minutes in a 37°C incubator), for a total of 6 freeze-thaw cycles. Lysates were then centrifuged at 4°C for 20 minutes 13,300 g. Supernatants were removed and flash frozen on dry ice and stored at −80°C until use.

#### Western blotting

TC lysates were prepared for immunoblotting by mixing protein lysates with lammeli sample buffer (BioRad, cat# 1610747) and beta-mercaptoethanol (BioRad, cat# 1610710) for a final concentration of 30 μg/mL total protein, and boiling at 95°C for 5 minutes. Proteins were separated using gel electrophoresis with hand-cast SDS-page gels (4% stacking gel [0.5M Tris-HCl Stacking buffer, BioRad cat# 1610799], 8% resolving gel [1.5M Tris-HCl Resolving buffer], 10% APS, 10% SDS, TEMED (Invitrogen, ref#15524-10) or BioRad 4-20% mini-PROTEAN TGX gels (BioRad, cat#4561091) using the BioRad mini-PROTEAN system. The Biorad Precision Plus Kaleidoscope ladder (Biorad, cat# 1610375) was used as a molecular weight standard. After rapid semi-dry transfer to a nitrocellulose membrane (Trans-Blot Turbo Transfer, Bio-Rad, cat#1704159), Ponceau S staining solution (Thermo Scientific, cat#A40000279) was used to evaluate transfer efficiency and visualize protein bands for total protein normalization. Membranes were blocked in 3% milk in TBST for 1 hour at room temperature before antibody application. Primary antibodies were diluted in 3% BSA in TBST and incubated at 4°C overnight for 16-20 hours on an orbital shaker. The following day, membranes were washed in TBST and incubated with a HRP-conjugated secondary antibody diluted in 3% milk in TBST for 2 hours at room temperature on an orbital shaker. Membranes were incubated with SuperSignal West Pico PLUS Chemiluminescent Substrate (ThermoFisher, cat#34095) for 5 minutes before imaging on Azure Biosystems Ai600 Imager.

#### Densitometry

Full-sized images of western blots are included in supplemental data. Densitometry was used to quantitate western blot images following the total protein normalization procedure outlined by Hossein Davarinejad (www.yorku.ca/yisheng/Internal/Protocols/ImageJ.pdf). Protein band intensity was subtracted from background membrane intensity to obtain net band intensity, which was normalized to full-lane Ponceau intensity to get a ratio of net protein/net Ponceau.

#### Microscopy & Image analysis

Images were taken with an AxioImager Z2 with apotome correction (Zeiss). Neurons were imaged at 40x. For LTD stimulation, images were processed with deconvolution (fast iterative) and rolling ball background subtraction (diameter = 100). Intensity was calculated in Zen 3.9 (Zeiss).

### Proteomics

#### Sample processing

Proteomic data was acquired by the GUMC-Proteomics Shared Resource. Proteomic analysis was done following an established protocol [20, 22, 23]. Briefly, hippocampal lysates were aliquoted to 100 μg/mL of total protein and were loaded into an S-trap column (ProtiFi, LCC) for digestion (trypsin/Lys-C). Digests were eluted and dried using a SpeedVac (Fisher Scientific). Nano ultra-performance liquid chromatography coupled to tandem mass spectrometry (UPLC-MS/MS; nanoAcquity UPLC coupled with Orbitrap Fusion Lumos mass spectrometer) was used to separate and analyze peptides. Data was acquired in data-independent acquisition (DIA) mode (ion spray voltage = 2.4 kV; ion transfer temperature = 275°C). Mass spectra were recorded using the Xcalibur 4.0 with advanced peak determination for MS analyses.

#### Proteomic analyses

Proteomic results were analyzed using the Spectronaut software (Biognosys, version 19) in DirectDIA (default settings). The Qvalve Cutoff was set to 1% at peptide precursor protein level (scrambled decoy generation, dynamic size = 0.1 fraction of library size). MS2-based quantification by area was used to enable local cross-run normalization. Results were then fed into a logistic Bayesian model that compared the linear fit of each protein to a training dataset comprised of data collected for each animal, compared to one outcome variable, i.e. genotype. Briefly, data was fitted to multiple linear models representing training data. Resulting data was log-transformed and fitted as a function of genotype to each training variable. Hypothesis testing (two-tailed, 5% alpha) for genotype prediction were used in each model. Model fit, visualization, and GO enrichment analysis were done in R (2024).

### Statistical analyses

All hypothesis testing was done using Prism (GraphPad, Version 10.4.1). All data was tested for normality (Shapiro-Wilk test) and skew (D’Agostino-Pearson omnibus (K2)), and normalized (log-transform; z-score) as needed. Outlier analysis (ROUT) was used to remove outliers. Effect size (Cohen’s *d*) was calculated using G*Power (Version 3.1.9.6). T-tests (two-tailed, α = 0.05) are graphed showing standard error of the mean (SEM). Analysis of variance, including ANOVA, 2-way ANOVA, and mixed-effects analysis (REML) (α = 0.05), were followed by post-hoc multiple comparisons (Tukey’s HSD) and the *q*-value (adjusted *p*-value) was used to determine statistical significance (* = *p* < 0.05, ** = *p* < 0.005, *** = *p* < 0.0005, **** = *p* < 0.0001). Error bars for all graphs show the SEM.

All code generated for EEG and proteomic analysis can be found at: https://github.com/KatHummel/CCR5.

## Acknowledgments

We would like to acknowledge the Division of Comparative Medicine for their veterinary support, as well as Dr. Sung Hyeok Hong for his assistance with telemeter surgeries.

## References

1. Thompson, S.M., et al., An excitatory synapse hypothesis of depression. Trends Neurosci, 2015. 38(5): p. 279–94.

2. Duman, R.S. and N. Li, A neurotrophic hypothesis of depression: role of synaptogenesis in the actions of NMDA receptor antagonists. Philos Trans R Soc Lond B Biol Sci, 2012. 367(1601): p. 2475–84.

3. Duman, R.S. and L.M. Monteggia, A neurotrophic model for stress-related mood disorders. Biol Psychiatry, 2006. 59(12): p. 1116–27.

4. Nestler, E.J., et al., Neurobiology of depression. Neuron, 2002. 34(1): p. 13–25.

5. Alaiyed, S., et al., Venlafaxine Stimulates an MMP-9-Dependent Increase in Excitatory/Inhibitory Balance in a Stress Model of Depression. J Neurosci, 2020. 40(22): p. 4418–4431.

6. Riga, D., et al., Hippocampal extracellular matrix alterations contribute to cognitive impairment associated with a chronic depressive-like state in rats. Sci Transl Med, 2017. 9(421).

7. Gonzalez, E.O., et al., The effects of Maraviroc on liver fibrosis in HIV/HCV co-infected patients. J Int AIDS Soc, 2014. 17(4 Suppl 3): p. 19643.

8. Lefebvre, E., et al., Antifibrotic Effects of the Dual CCR2/CCR5 Antagonist Cenicriviroc in Animal Models of Liver and Kidney Fibrosis. PLoS One, 2016. 11(6): p. e0158156.

9. Ulbrich, P., et al., Interplay between perivascular and perineuronal extracellular matrix remodelling in neurological and psychiatric diseases. Eur J Neurosci, 2021. 53(12): p. 3811–3830.

10. Bauer, O., et al., Association of Chemokine (C-C Motif) Receptor 5 and Ligand 5 with Recovery from Major Depressive Disorder and Related Neurocognitive Impairment. Neuroimmunomodulation, 2020. 27(3): p. 152–162.

11. Oglodek, E.A., et al., Comparison of chemokines (CCL-5 and SDF-1), chemokine receptors (CCR-5 and CXCR-4) and IL-6 levels in patients with different severities of depression. Pharmacol Rep, 2014. 66(5): p. 920–6.

12. Yao, H., et al., Astrocyte-derived CCL5-mediated CCR5(+) neutrophil infiltration drives depression pathogenesis. Sci Adv, 2025. 11(21): p. eadt6632.

13. Zhou, M., et al., CCR5 is a suppressor for cortical plasticity and hippocampal learning and memory. Elife, 2016. 5.

14. Khalid, A., et al., Gamma oscillation in functional brain networks is involved in the spontaneous remission of depressive behavior induced by chronic restraint stress in mice. BMC Neurosci, 2016. 17: p. 4.

15. Arikan, M.K., B. Metin, and N. Tarhan, EEG gamma synchronization is associated with response to paroxetine treatment. J Affect Disord, 2018. 235: p. 114–116.

16. Fitzgerald, P.J. and B.O. Watson, Gamma oscillations as a biomarker for major depression: an emerging topic. Transl Psychiatry, 2018. 8(1): p. 177.

17. Jetsonen, E., et al., Activation of TrkB in Parvalbumin interneurons is required for the promotion of reversal learning in spatial and fear memory by antidepressants. Neuropsychopharmacology, 2023. 48(7): p. 1021–1030.

18. Alaiyed, S., et al., Venlafaxine stimulates PNN proteolysis and MMP-9-dependent enhancement of gamma power; relevance to antidepressant efficacy. J Neurochem, 2019. 148(6): p. 810–821.

19. Greco, G.A., et al., CCR5 deficiency normalizes TIMP levels, working memory, and gamma oscillation power in APOE4 targeted replacement mice. Neurobiol Dis, 2023. 179: p. 106057.

20. Gilbert, K.F., et al., Pramipexole, a D3 receptor agonist, increases cortical gamma power and biochemical correlates of cortical excitation; implications for mood disorders. Eur J Neurosci, 2024.

21. Lee, J. and D.T.S. Pak, Amyloid precursor protein combinatorial phosphorylation code regulates AMPA receptor removal during distinct forms of synaptic plasticity. Biochem Biophys Res Commun, 2024. 709: p. 149803.

22. Bauman, T.A., et al., Serum fibrinogen is not elevated in patients with myasthenia gravis. Sci Rep, 2025. 15(1): p. 13013.

23. Wu, C., et al., Coupling suspension trapping-based sample preparation and dataindependent acquisition mass spectrometry for sensitive exosomal proteomic analysis. Anal Bioanal Chem, 2022. 414(8): p. 2585–2595.

24. Buzsaki, G., et al., Pattern and inhibition-dependent invasion of pyramidal cell dendrites by fast spikes in the hippocampus in vivo. Proc Natl Acad Sci U S A, 1996. 93(18): p. 9921–5.

25. Gartner, M., et al., Aberrant working memory processing in major depression: evidence from multivoxel pattern classification. Neuropsychopharmacology, 2018. 43(9): p. 1972–1979.

26. Povsic, T.J., T.A. Kohout, and R.J. Lefkowitz, Beta-arrestin1 mediates insulin-like growth factor 1 (IGF-1) activation of phosphatidylinositol 3-kinase (PI3K) and anti-apoptosis. J Biol Chem, 2003. 278(51): p. 51334–9.

27. Peineau, S., et al., A systematic investigation of the protein kinases involved in NMDA receptor-dependent LTD: evidence for a role of GSK-3 but not other serine/threonine kinases. Mol Brain, 2009. 2: p. 22.

28. Dong, Z., et al., Hippocampal long-term depression mediates spatial reversal learning in the Morris water maze. Neuropharmacology, 2013. 64: p. 65–73.

29. Kim, J.I., et al., PI3Kgamma is required for NMDA receptor-dependent long-term depression and behavioral flexibility. Nat Neurosci, 2011. 14(11): p. 1447–54.

30. Struyf, S., et al., Natural truncation of RANTES abolishes signaling through the CC chemokine receptors CCR1 and CCR3, impairs its chemotactic potency and generates a CC chemokine inhibitor. Eur J Immunol, 1998. 28(4): p. 1262–71.

31. Monaco, S.A., A.J. Matamoros, and W.J. Gao, Conditional GSK3beta deletion in parvalbumin-expressing interneurons potentiates excitatory synaptic function and learning in adult mice. Prog Neuropsychopharmacol Biol Psychiatry, 2020. 100: p. 109901.

32. Tewari, B.P., et al., Perineuronal nets decrease membrane capacitance of peritumoral fast spiking interneurons in a model of epilepsy. Nat Commun, 2018. 9(1): p. 4724.

33. Lensjo, K.K., et al., Removal of Perineuronal Nets Unlocks Juvenile Plasticity Through Network Mechanisms of Decreased Inhibition and Increased Gamma Activity. J Neurosci, 2017. 37(5): p. 1269–1283.

34. Gurevicius, K., et al., Genetic ablation of tenascin-C expression leads to abnormal hippocampal CA1 structure and electrical activity in vivo. Hippocampus, 2009. 19(12): p. 1232–46.

35. Happel, M.F., et al., Enhanced cognitive flexibility in reversal learning induced by removal of the extracellular matrix in auditory cortex. Proc Natl Acad Sci U S A, 2014. 111(7): p. 2800–5.

36. Kammel, L.G., et al., Enhanced GABAergic Tonic Inhibition Reduces Intrinsic Excitability of Hippocampal CA1 Pyramidal Cells in Experimental Autoimmune Encephalomyelitis. Neuroscience, 2018. 395: p. 89–100.

37. Wu, Z., et al., Tonic inhibition in dentate gyrus impairs long-term potentiation and memory in an Alzheimer’s [corrected] disease model. Nat Commun, 2014. 5: p. 4159.

38. Li, T. and J. Zhu, Entanglement of CCR5 and Alzheimer’s Disease. Front Aging Neurosci, 2019. 11: p. 209.

39. Wee, J.J. and S. Kumar, Prediction of hub genes of Alzheimer’s disease using a protein interaction network and functional enrichment analysis. Genomics Inform, 2020. 18(4): p. e39.

40. Kelder, W., et al., Beta-chemokines MCP-1 and RANTES are selectively increased in cerebrospinal fluid of patients with human immunodeficiency virus-associated dementia. Ann Neurol, 1998. 44(5): p. 831–5.

41. Hassamal, S., Chronic stress, neuroinflammation, and depression: an overview of pathophysiological mechanisms and emerging anti-inflammatories. Front Psychiatry, 2023. 14: p. 1130989.

42. Shen, Y., et al., CCR5 closes the temporal window for memory linking. Nature, 2022. 606(7912): p. 146–152.

43. Tene, O., et al., CCR5-Delta32 polymorphism: a possible protective factor for post-stroke depressive symptoms. J Psychiatry Neurosci, 2021. 46(4): p. E431–E440.

44. Zhang, W., et al., Altered fronto-central theta-gamma coupling in major depressive disorder during auditory steady-state responses. Clin Neurophysiol, 2023. 146: p. 65–76.

45. Singh, B., et al., Brain-wide human oscillatory local field potential activity during visual working memory. iScience, 2024. 27(3): p. 109130.

46. Khalaf, A., et al., Early neural activity changes associated with stimulus detection during visual conscious perception. Cereb Cortex, 2023. 33(4): p. 1347–1360.

47. Frame, S. and P. Cohen, GSK3 takes centre stage more than 20 years after its discovery. Biochem J, 2001. 359(Pt 1): p. 1–16.

48. Beaulieu, J.M., et al., A beta-arrestin 2 signaling complex mediates lithium action on behavior. Cell, 2008. 132(1): p. 125–36.

49. Lee, Y., et al., The GSK-3 Inhibitor CT99021 Enhances the Acquisition of Spatial Learning and the Accuracy of Spatial Memory. Front Mol Neurosci, 2021. 14: p. 804130.

50. Miranda, J.M., et al., Hippocampal parvalbumin interneurons play a critical role in memory development. Cell Rep, 2022. 41(7): p. 111643.

51. Wang, W., et al., Correlation of Electrophysiological and Gene Transcriptional Dysfunctions in Single Cortical Parvalbumin Neurons After Noise Trauma. Neuroscience, 2022. 482: p. 87–99.

52. de Villers-Sidani, E., et al., Manipulating critical period closure across different sectors of the primary auditory cortex. Nat Neurosci, 2008. 11(8): p. 957–65.

53. Sampedro-Piquero, P., et al., Environmental enrichment as a therapeutic avenue for anxiety in aged Wistar rats: Effect on cat odor exposition and GABAergic interneurons. Neuroscience, 2016. 330: p. 17–25.

54. Rowlands, D., et al., Aggrecan Directs Extracellular Matrix-Mediated Neuronal Plasticity. J Neurosci, 2018. 38(47): p. 10102–10113.

55. Lupori, L., et al., A comprehensive atlas of perineuronal net distribution and colocalization with parvalbumin in the adult mouse brain. Cell Rep, 2023. 42(7): p. 112788.

56. Winkel, F., et al., Pharmacological and optical activation of TrkB in Parvalbumin interneurons regulate intrinsic states to orchestrate cortical plasticity. Mol Psychiatry, 2021. 26(12): p. 7247–7256.

57. Aguilar, J.S. and A.W. Lasek, Modulation of stress-, pain-, and alcohol-related behaviors by perineuronal nets. Neurobiol Stress, 2024. 33: p. 100692.

58. Midwood, K., et al., Tenascin-C is an endogenous activator of Toll-like receptor 4 that is essential for maintaining inflammation in arthritic joint disease. Nat Med, 2009. 15(7): p. 774–80.

59. Gottschling, C., et al., Elimination of the four extracellular matrix molecules tenascin-C, tenascin-R, brevican and neurocan alters the ratio of excitatory and inhibitory synapses. Sci Rep, 2019. 9(1): p. 13939.

60. Bozzelli, P.L., et al., Proteolytic Remodeling of Perineuronal Nets: Effects on Synaptic Plasticity and Neuronal Population Dynamics. Neural Plast, 2018. 2018: p. 5735789.

61. Koshikawa, Y., et al., Disentangling cognitive inflexibility in major depressive disorder: A transcranial direct current stimulation study. Psychiatry Clin Neurosci, 2022. 76(7): p. 329–337.

62. Provenzano, J., et al., Inflexibly sustained negative affect and rumination independently link default mode network efficiency to subclinical depressive symptoms. J Affect Disord, 2021. 293: p. 347–354.

